# Integrated transcriptomic and methylome analysis reveals retinoic acid pathway activation after decitabine treatment in EBV associated gastric cancer

**DOI:** 10.1101/2025.10.14.682272

**Authors:** Sarah Preston-Alp, Yin Wang, Matteo Bergonzoni, Andrew Kossenkov, Samantha Soldan, Lisa Beatrice Caruso, Davide Maestri, Paul M. Lieberman, Benjamin Gewurz, Italo Tempera

## Abstract

Epstein-Barr virus associated gastric cancer (EBVaGC) accounts for ∼9–10% of gastric cancers worldwide and is defined by a distinctive molecular profile, including extreme hypermethylation of the DNA. Targeting this aberrant methylation may be a potential therapeutic strategy. EBV^+^ gastric cancer cell lines (YCCEL1, SNU719) and EBV^-^ lines (AGS, SNU16, MKN74) were treated with a DNA methyl-transferase inhibitor (DNMT), decitabine (DCB), for three days followed by RNA sequencing to identify EBV-specific responses. DNA methylation profiling by reduced representation bisulfite sequencing (RRBS) was performed in EBV^+^ cell-lines and integrated with expression data to identify epigenetically regulated networks. While DCB induced broad transcriptional changes across all lines, EBV^+^ cells exhibited the strongest transcriptional response, sharing many upregulated genes. Many of these EBV^+^ specific genes were expressed at lower baseline levels in EBV^+^ tumors from TCGA. DCB predominantly reduced methylation at highly methylated intergenic CpGs, with a subset of promoters undergoing significant demethylation. Integrated analysis revealed a strong inverse correlation between promoter demethylation and gene expression, implicating multiple cancer-relevant pathways. Upstream regulator analysis and motif enrichment indicated that regions losing methylation were enriched for retinoic acid receptor α (RARα) binding motifs, suggesting that DCB-mediated demethylation restores RA pathway accessibility and transcriptional activity. Further, inhibiting RAR signaling reduced DCB induced apoptosis. Although DCB can induce both host gene re-expression and viral lytic gene activation in EBV-positive tumors, its impact on RA signaling in EBVaGC has not been studied. Decitabine promotes extensive epigenetic reprogramming in EBVaGC, with preferential effects in CIMP-positive, EBV-infected cell lines.

**Importance:** EBV^+^ gastric cancer contains hypermethylated DNA and despite this distinct molecular phenotype there are currently no EBV-specific treatments available. Using an FDA approved inhibitor to target hypermethylated DNA and multi-omics approach to study the cellular response, we uncovered epigenetically altered transcriptional networks that may be further exploited to improve potential therapy. Among the pathways disrupted, retinoic acid signaling is of particular interest, as retinoid receptors such as RARα and RARβ are frequently hypermethylated and repressed in EBVaGC. Our findings indicate that DNMT inhibition can partially reverse RA receptor silencing, supporting further investigation of DNMTi–RA combination strategies as a novel therapy for EBV^+^ gastric cancer.

## Introduction

Epstein–Barr virus (EBV) is a ubiquitous herpesvirus that infects more than 90% of adults worldwide and drives multiple human malignancies (e.g. Burkitt lymphoma, nasopharyngeal carcinoma), including a distinct subset of gastric carcinomas. EBV positive gastric cancer (EBVaGC) accounts for 9-10% of gastric cancers worldwide and display distinct molecular characteristics[1-3]. EBVaGC tends to occur in relatively younger patients with a male predominance, often arising in the proximal stomach and characterized by lymphocytic infiltration in the tumor [4, 5]. The median survival of EBVaGC is 8.5 years with a better prognosis than EBV negative cases[4, 6]. Molecularly, EBVaGC display extensive widespread DNA hypermethylation and extreme CpG island methylator phenotype (CIMP), with a markedly high frequency of promoter CpG island methylation and transcriptional silencing of many tumor suppressor and differentiation genes [7-10]. EBV infection appears to drive DNA hypermethylation as both the host and viral genomes are hypermethylated after gastric epithelial cell infection of EBV [11].

DNA methylation (5mC) is catalyzed by the DNA methyltransferase (DNMT) enzyme family by placing a methyl group on the 5-carbon of cytosine in cytosine-guanine dinucleotide pairs. DNMT3A/B are responsible for de novo 5mC while DNMT1 maintains 5mC patterning during cell division. Pharmacologic inhibition of DNMTs with a cytidine analogue, decitabine (5-aza-2’-deoxycytidine, DCB) induces passive loss of 5mC through cell divisions leading to expression of silenced host genes in both solid and hematological malignancies[12-14]. In gastric cancer, DCB treatment has been shown to enhance cellular differentiation[15, 16]. In EBVaGC, DCB treatment extensively hypomethylated the EBV episome and induced expression of lytic-phase viral genes [17, 18] . Our previous study showed that DCB disorganizes methylation patterning around critical CTCF boundaries necessary to maintain the episomal 3D-architecture, to increase chromatin accessibility and promote lytic-gene expression [17, 19]. However, the downstream molecular pathways reactivated within the host signaling network by decitabine in EBVaGC and how they contribute to tumor control have not been fully examined.

One pathway of particular interest is the retinoic acid (RA) pathway. All-trans retinoic acid (ATRA), the bioactive metabolite of vitamin A, binds nuclear retinoid receptors, RAR*α*, RAR*β*, and RAR*γ* which dimerize to engage retinoic acid response elements within the DNA sequences to control cell differentiation and growth arrest [20]. The use of ATRA as a differentiation therapy in acute promyelocytic leukemia established retinoic acid signaling as a powerful anti-cancer mechanism when appropriately engaged [21, 22]. In solid tumors, retinoic acid maintains epithelial cell differentiation and proliferation with loss of RA signaling frequently associated with tumor development and progression [23-25]. In various cancers, including gastric cancer, RAR*β* gene expression is silenced through extensive promoter methylation in approximately 64% of gastric cancers which has been linked to worse prognosis and aggressive disease [26]. In the context of EBV-related cancers, retinoic acid signaling appears to be especially relevant and intriguingly disrupted. While the lytic EBV protein BZLF1 appears to activate RA signaling, EBV’s latent oncoprotein LMP1 can actively interfere with the RA pathway by upregulating host DNMTs and drive hypermethylation of the *RARβ2* gene promoter [27-29]. Such findings illustrate that EBV can modulate host nuclear receptor pathways to favor oncogenesis. Consistent with this, comprehensive methylome analyses of EBV-associated gastric tumors have identified RA pathway genes as frequent targets of viral-induced epigenetic silencing. For example, RARA was found among the pro-differentiation genes recurrently hypermethylated in EBV-positive gastric cancers [30]. The epigenetic inactivation of RARα, a key mediator of retinoid signaling, further suggests that RA pathways are suppressed or altered in EBVaGC as part of its oncogenic program. Taken together, the current evidence indicates that EBVaGC harbor a RA pathway that is likely dormant due to the virus’s epigenetic influence.

To understand the effects of DCB on GC, we performed transcriptomic analysis on a panel of EBV positive and negative cell lines. Transcriptomic analysis identified general effects induced by DCB across cell lines, as well as possible, CIMP specific and EBV specific transcriptional responses. We performed Reduced Representation Bisulfite Sequencing (RRBS) to analyze the changes in 5mC across the host genome within EBV+ cells and found epigenetic manipulation of RA among several others. Our data suggest DCB treatment may restore RA and its growth inhibitory effects.

## Materials and methods

### Cell Culture and treatments

This study used EBV^+^ cell lines SNU719 and YCCEL1, and EBV-cell lines AGS, MKN74, and SNU-16 [31, 32]. Cell lines were cultured in RPMI 1640 medium supplemented with 10% FBS at 37°C with 5% CO2. For retinoic acid treatments, cells were first treated 3d with 1µM DCB followed by 3 and 6d treatment of all-trans retinoic acid or AR-7 alone or in combination with DCB. For PI/Annexin V flow cytometry 10^5^ cells were harvested using trypsin, washed twice in cold PBS and resuspended in 100µL Annexin V Binding Buffer. Cells were stained with 5*μ*L AnnexinV-FITC and 5*μ*L propidium iodide (BioLegend) and incubated for 15min at room temperature before adding 100*μ*L binding buffer and analyzing on the flow cytometer. Significance was calculated using a one-way ANOVA followed by a Tukey’s HSD post-hoc test.

#### RNA-sequencing

RNA was isolated and sequenced as previously described[17]. RNA was extracted using the RNeasy Mini Kit (Qiagen) with on column DNase I digestion, according to the manufacturers protocol. RNA libraries were generated using the QuantSeq (Lexogen) library prep by oligo-dT priming to produce strand-specific libraries and were sequenced on the NextSeq500 (Illumina) to generate single-end 76bp reads [33].

### Reduced Representation Bisulfite Sequencing – RRBS

DNA was extracted using ThermoFisher GeneJet Genomic DNA Isolation kit according to the manufacturers protocol. Digestions were performed with 2*μ*g of DNA with 100 U of MspI endonuclease at 37°C for 4h. End filling and 3’dA overhangs were created using Klenow fragment (3’>5’ exo-) DNA polymerase. Reactions were cleaned using 2x AMPure XP bead solution. T4 DNA ligase was used to ligated methylated adapters to the DNA fragments at 16°C overnight. Adapters were cleaved with USER enzyme at 37°C for 15 min. Dual size selection was performed using AMPure XP beads to maintain DNA fragments between 150-500bp. Purified fragments were bisulfite converted using the KIT according to the manufacturers protocol. The bisulfite-converted fragments were cleaned using 3x AMPure XP bead volume. Fragments were indexed using the NEBNext Index i7 primers for 15 cycles. Libraries were pooled to a concentration of 10nM and spiked 25% with phiX to increase sequence diversity.

### Bioinformatics

RNA-seq reads were aligned to the human reference genome (GRCh38) and the EBV genome (NC_007605_1) using STAR [34]. Raw counts were analyzed with DESeq2 including only genes with greater than 10 counts across samples. Wald’s test was used to compute p-value between contrasts and corrected for multiple testing using Benjamini-Hochberg[35]. Log2 fold-changes were adjusted using DESeq2 apeglm. Differentially expressed genes were called with greater than absolute log2 Fold Change > 2 and q<0.05. RRBS reads were trimmed with adapter and M-bias-trimming using Cutadapt with –rrbs and default quality filtering. Trimmed reads were aligned with Bismark [36, 37]. Methylation calls were generated with Bismark_methylation_extractor. Cytosine coverage files were used for downstream analyses. CpG methylation was calculated in R from Bismark coverage files. CpG sites were filtered for coverage greater than 20 reads. Common CpG sites across samples were unified with methyKit unite [38]. Differential methylation was tested with calculateDiffMeth and corrected for overdispersion; FDR was corrected using Benjamini-Hochberg. Promoter CpG sites were defined as -1000 to 0 bp upstream of the transcriptional start site. Known transcription factor motif enrichment was performed with HOMER [39]. Differentially methylated CpG sites were expanded to 50bp windows centered on the CpG and background regions were GC- and length-matched using all RRBS-detectable CpGs that did not change significantly, to control for RRBS selection bias. Enrichment was calculated using HOMER’s hypergeometric test with Benjamini-Hochberg corrections; significance was considered with q < 0.05. Data was accessed under accession number GSE239770 and GSE234658.

## Results

### DCB induces global transcriptional upregulation of gastric cancer cells

To understand the effect of DNMT inhibition by small molecule inhibition using Decitabine, gastric cancer cell lines YCCEL1, SNU 719, AGS, SNU-16, and MKN-74 were treated with 7.5uM DCB for 3d and transcriptomic profiling by RNA-sequencing was performed. Principal component analysis of variance on significantly changed genes shows concordant changes by treatment along PC1 along with contribution based on cell type along PC2 (Figure 1A). EBV positive cell lines, YCCEL1 and SNU-719 are shown clustered together indicating transcriptional similarity both basally and in response to DCB. Overall, DCB lead to an increase in gene expression of 820 genes with few genes being significantly down-regulated consistent with the idea that methylation loss allows for transcriptional activation (Figure 1B). Further, a heatmap of all differentially expressed genes shows all CIMP^+^ cells (AGS, SNU719, and YCCEL1) clustering together before treatment but EBV^-^ AGS cell line more like CIMP^-^ cells (HGC27 and SNU16) after DCB treatment (Figure 1C). Gene set enrichment analysis of these DCB effected genes show changes in pathways related to neuronal pathogenesis and solid cancers (Figure 1D). Overall, we find a variable effect in transcriptional response among the cell lines profiled with the EBV^+^ and CIMP^+^ cells being most effected (Figure 1E).

**Figure 1.**
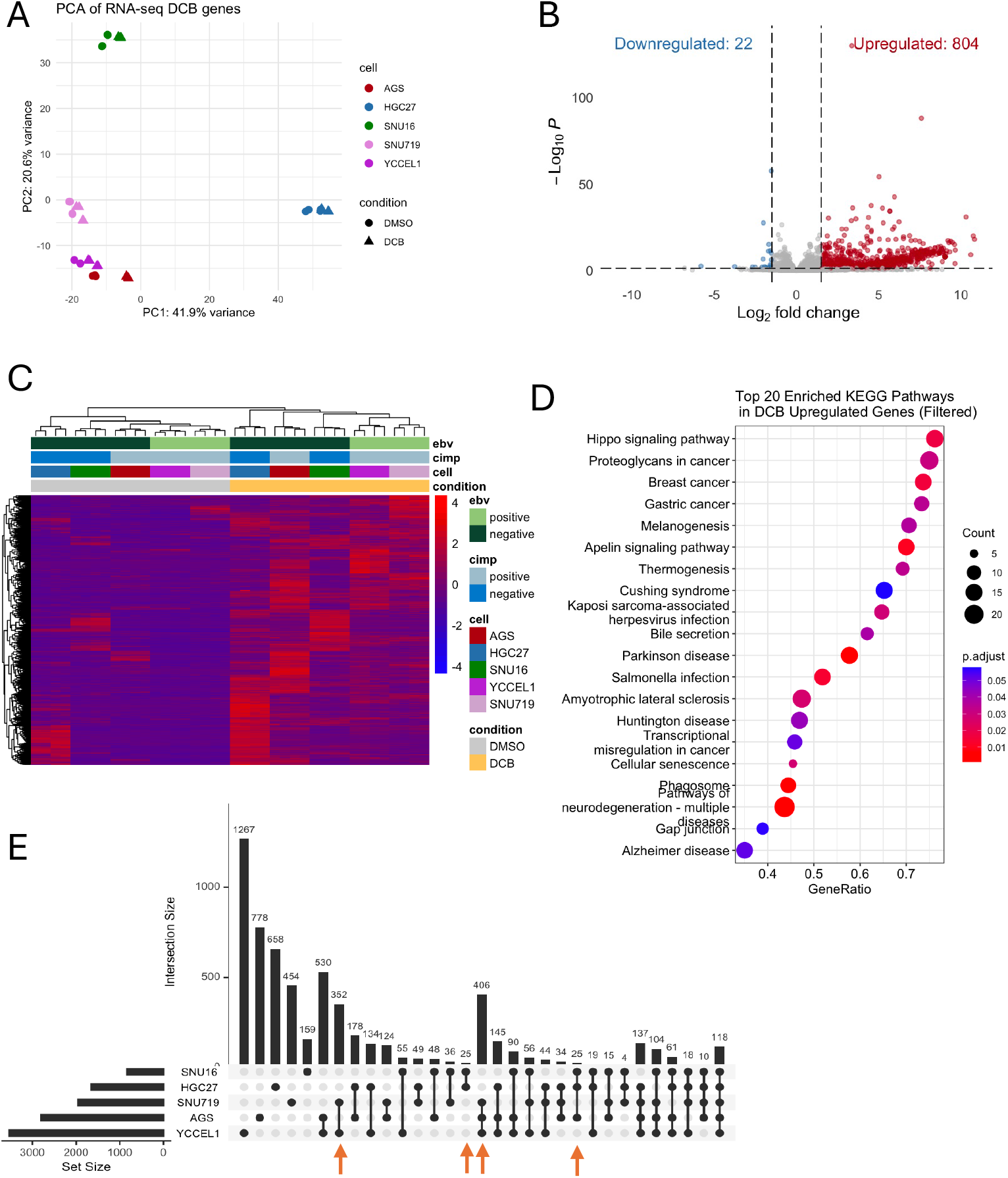
DCB induces global transcriptional upregulation of gastric cancer cells. A) Principal component analysis of differentially expressed genes after 3d of DMSO or 7.5 *μ*M decitabine treatment for gastric cancer cell lines. B) Volcano plot of gene expression after DCB treatment showing upregulated genes in red and down regulated-genes in blue. C) Heatmap of differentially expressed genes (log_2_ fold change > 2 & adjusted *P* < 0.05) after DCB treatment. D) Enrichment analysis of differentially upregulated genes showing the top 20 KEGG pathways. E) An upset plot showing differentially expressed genes by cell-line after DCB treatment. Highlighted, are EBV-specific changes by the blue arrow and CIMP-specific changes by the green arrow.

### Distinct Gene Expression and Pathway Responses to DNA Demethylation in CIMP+and EBV+Gastric Cancer Cells

Since gastric cancer can showcase different phenotypes based on levels of DNA methylation either CIMP^+^ or CIMP^-^ we wanted to determine if there was a CIMP^+^-specific effect. We saw a high overlap between CIMP^+^ cell lines YCCEL1 and SNU719, and AGS with few unique gene expressions in MKN74 and SNU16. We found 406 genes upregulated in CIMP^+^ in response to DCB (Figure 2A). Gene ontology enrichment analysis of these genes identified over-representation of up-regulated genes in pathways related to intracellular transport, keratinocyte differentiation, and meiotic nuclear division. Pathways involving DNA methylation and apoptosis were also enriched. A cnet-plot was constructed to display how these enriched pathways and genes may be interconnected revealing distinct expression programs (Figure 2C). In one of the largest nodules involving DNA methylation, intermediate filament-based process, ncRNA metabolic, keratinocyte differentiation and epidermal cell differentiation process involve the upregulation of KRT family of proteins, MSX1, germline factor DDX4, mitotic checkpoint CHFR, and INHBB whose expression is associated with immune infiltrates in gastric cancers [40-42]. We further refined the analysis since CIMP^+^ GC can be either EBV^+^ or EBV^-^. We wanted to see if there were any EBV-specific effects, as we saw a large overlap in upregulated transcripts between YCCEL1 and SNU719 cells. We found 79 genes upregulated after DCB in EBV^+^ cells (Figure 3A). Some of these EBV-specific genes include *CDKN1C* which encodes *P57* and is a negative cell cycle regulator; *INPP4B* which is a PI3K pathway inhibitor; *NKD1* and *SFRP1* which are negative regulators of WNT signaling [43-48]. These genes are minimally expressed in the EBV^+^ cells where they show the greatest change in expression after DCB treatment compared to the EBV^-^ cell lines (Figure 3A,B). We compared the expression of these genes in the TCGA-STAUD cohort and found the EBV^+^ tumors had significantly lower expression of these EBV-specific DCB responsive genes than EBV^-^ negative tumors or normal tissue (Figure 3C). Gene ontology analysis of EBV^-^ specific response found an enrichment in inflammatory response, signal transduction, and cellular response to stimulus (Figure 3D). In line with this, DCB treatment reactivated immune related genes hypermethylated in EBVaGC, likely to help the virus evade the immune system [49].

**Figure 2.**
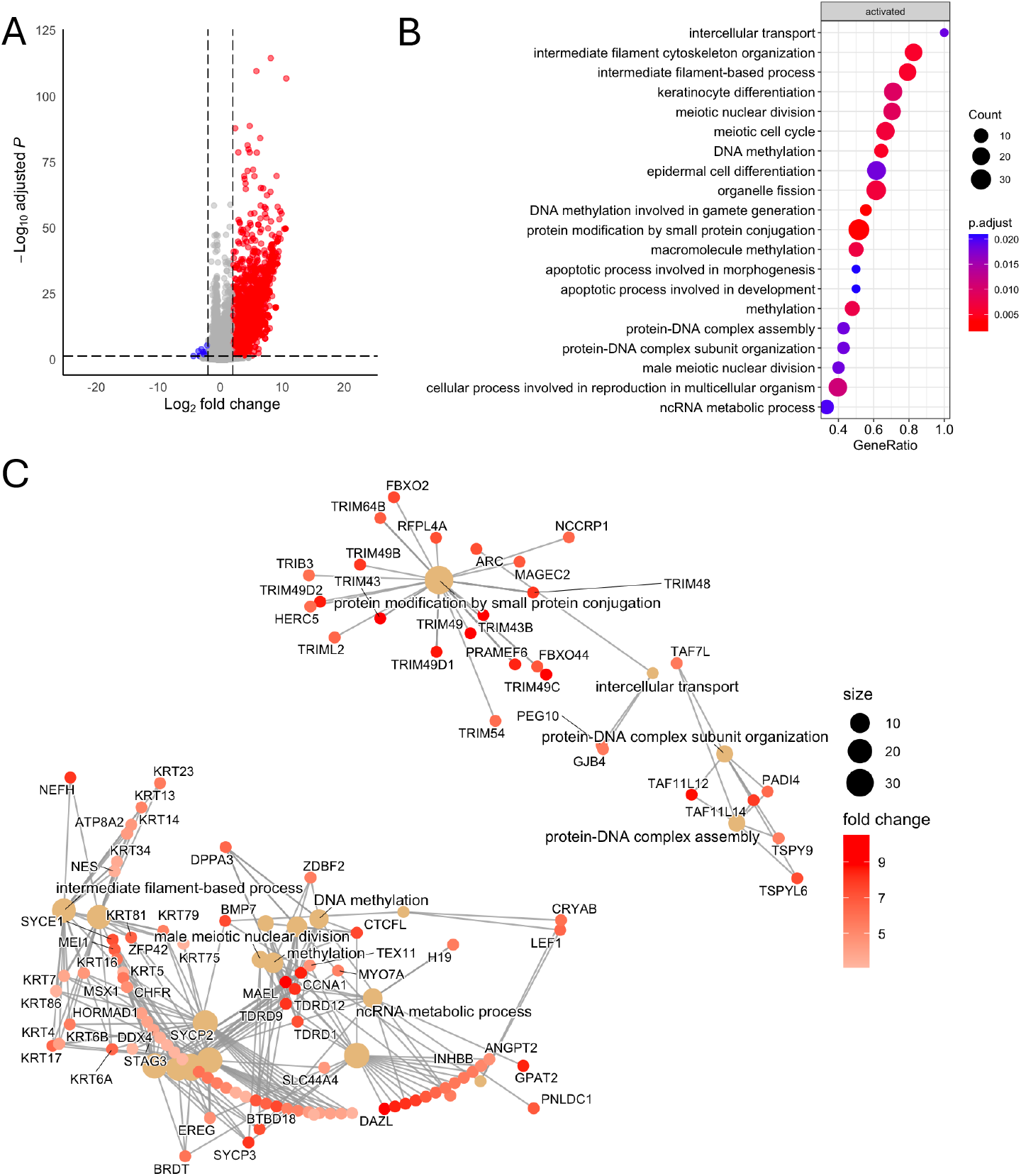
DCB transcriptional effects specific to cell lines that are hypermethylated. A) Volcano plot showing differentially expressed genes after DCB treatment in the hypermethylated cell lines: YCCEL1, SNU719, and AGS. B) GSEA analysis of DCB effect on CIMP^+^ cell lines C) Cnet plot of top 20 GSEA identified pathways and associated genes colored by fold change after DCB treatment.

**Figure 3.**
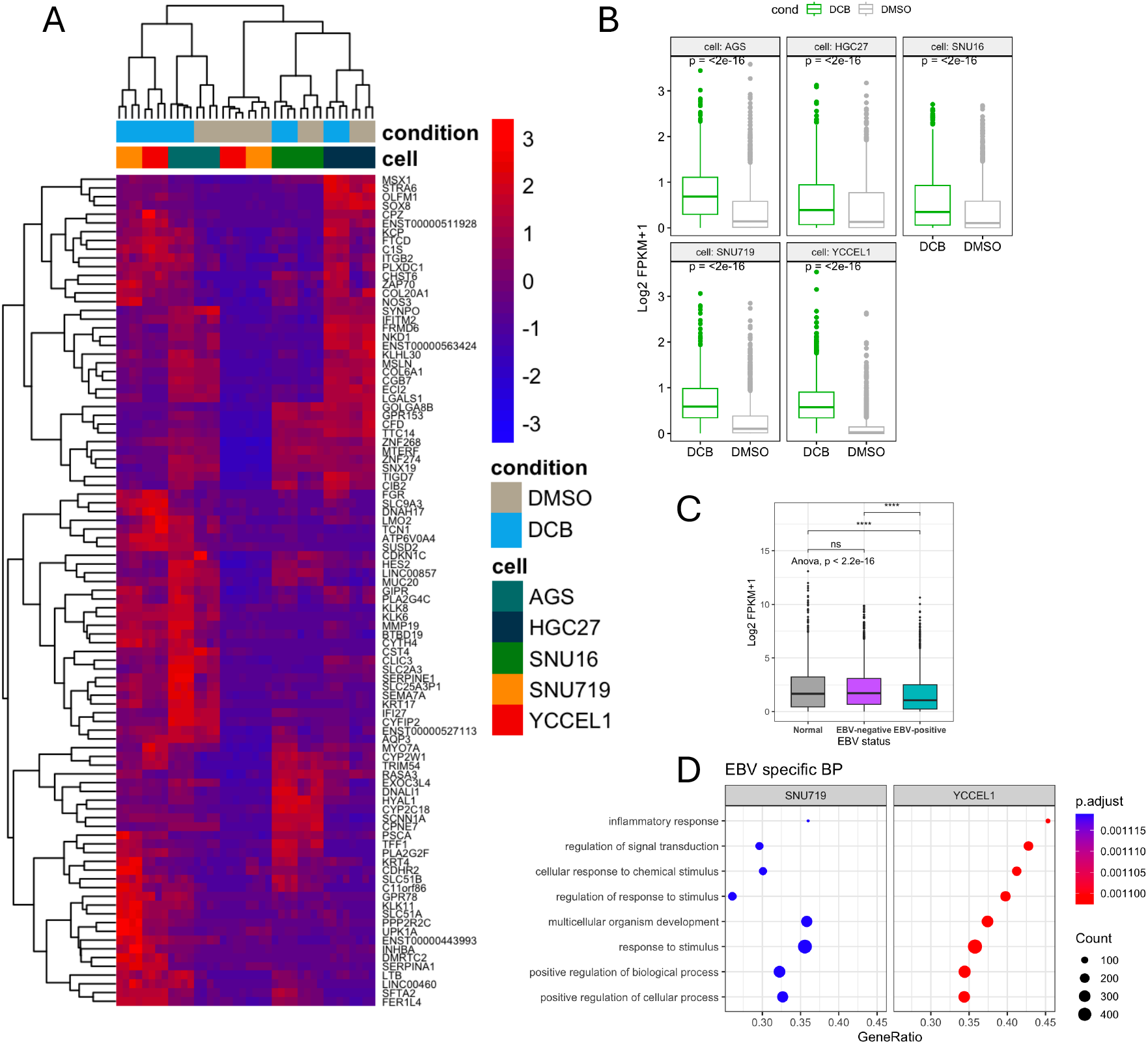
DCB transcriptional effects specific to EBV^+^ cell lines. A) Heatmap showing 79 EBV-specific differentially expressed genes (log_2_FC > 2) after DCB treatment. B) Violin plot of FPKM of EBV-specific genes across cell lines after DCB treatment. C) Violin plot of FPKM values of EBV-specific genes in the TCGA-STAD dataset separated by normal tissue or EBV status. D) GSEA of biological process analysis of DCB effect on EBV, showing only pathways specific for EBV.

### Genome-wide DNA Demethylation by DCB preferentially targets hypermethylated intergenic and intronic regions in EBV+gastric cancer cells

Since DNMTi by DCB leads to a global loss of DNA methylation allowing transcriptional activation of many genes, particularly in the EBV^+^ cell lines, we performed RRBS in YCCEL1 and SNU719 cell lines to understand the global impact of DCB induced 5mC changes on the transcriptional changes observed. PCA analysis on CpG 5mC showed a large separation between treatment by PC1 and cell line separation on PC2 (Figure 4A). Changes in DNA methylation occurred predominantly at hyper methylated CpG sites (>80%) leading to a shift towards an intermediate level of 5mC for both YCCEL1 and SNU719 (Figure 4B). In the SNU719 cell line we found 10% of the detectable CpG sites were hypomethylated by greater than 10% methylation after DCB treatment, compared to 5% of CpG sites in the YCCEL1 cell line (Figure 4C). A heatmap of significantly changed sites with greater than 20% change in methylation showed many changes commonly affected between cell lines on hypermethylated sites (Figure 4D). Many of the cell line specific effects were due to uniquely hypermethylated sites between the cell lines. Stacked bar plots display the proportion of all detectable CpG sites and their location within the promoter, intron, exon or intergenic region, compared to the proportion of hypomethylated sites (Figure 4E). In both cell lines, CpG sites within promoter regions were resistant to hypomethylation after DCB treatment whereas a larger proportion of CpG sites within intergenic and intronic regions were susceptible to hypomethylation.

**Figure 4.**
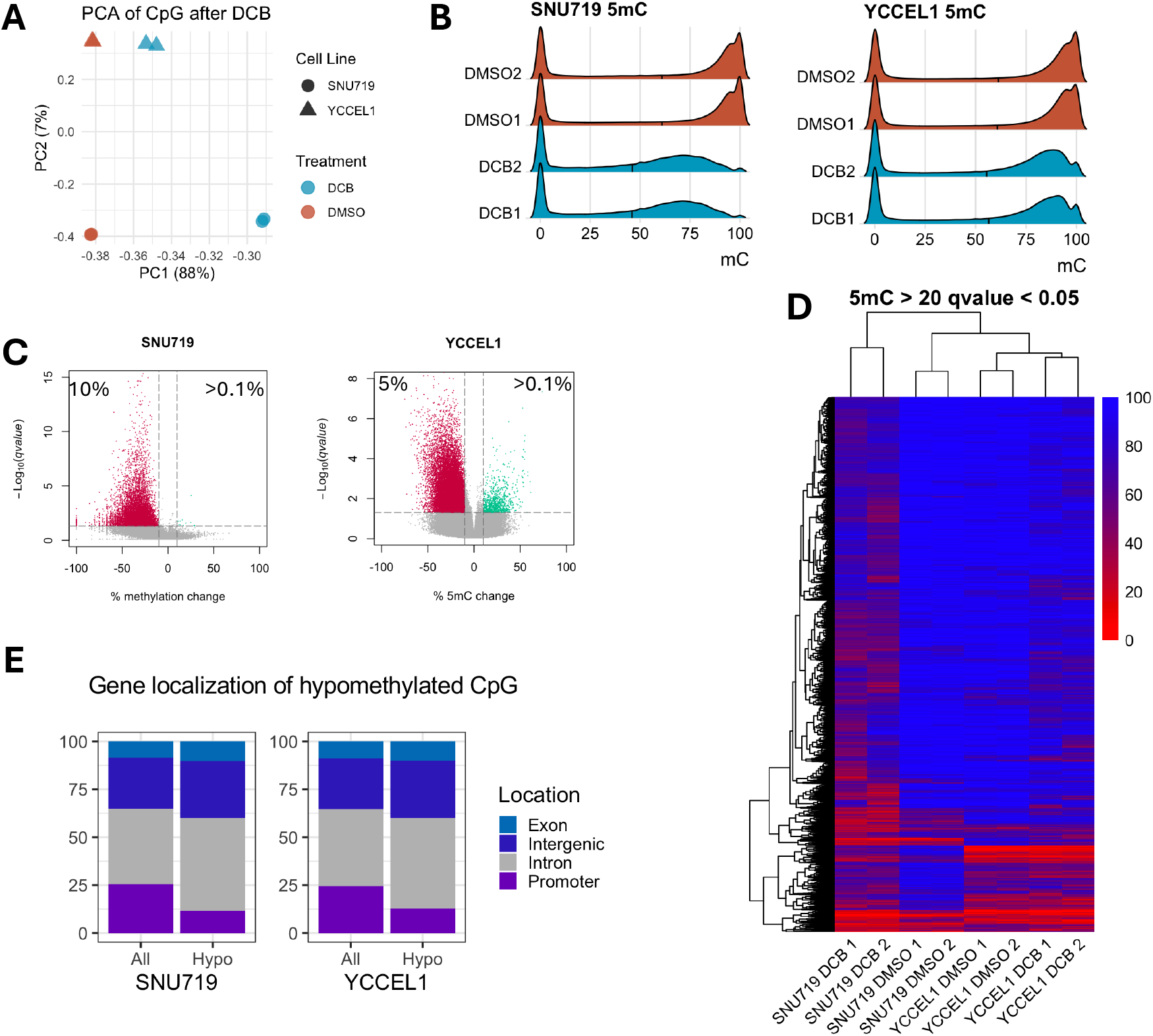
DCB transcriptional effects specific to EBV^+^ cell lines. A) PCA on CpG methylation from RRBS in SNU719 and YCCEL1 cell lines after DCB treatment. B) Ridgeline plot of CpG distribution in each sample. C) Volcano plot for each cell line for changes in CpG methylation after DCB, significantly hypomethylated CpG sites are red and hypermethylated sites are shown in green (>10% methylation change and q<0.01). D) Heatmap showing CpG methylation of significantly changed CpG sites. E) Stacked bar plot showing distribution of CpG sites within promoter, intron, intergenic, and exon regions of all detectable sites and those that are significantly hypomethylated.

### Promoter hypomethylation of tumor suppressor genes correlates with EBV-specific re-expression following DCB treatment

Despite promoters being resistant to change, we did observe many changes in hypomethylation throughout the genome and genome tracks for two tumor suppressor genes illustrate the correlation between promoter methylation and gene expression (Figure 5A). Previous reports identified promoter hypermethylation of the *RASSF1* promoter in EBV^+^ tumors and after EBV infection [29]. *RASSF1* showed loss of promoter methylation in SNU719 and not YCCEL1 correlating with increased expression only in SNU719 after DCB treatment (Figure 5A,B). We observed increased expression in the EBV^-^ cell lines, AGS and SNU16, after DCB treatment suggesting *RASSF1* re-expression is not determined by EBV status. *HOPX* has been shown to have tumor suppressive effects in several cancers including nasopharyngeal carcinoma and gastric cancer where promoter methylation was associated with tumor stage [50, 51]. *HOPX* showed loss of promoter methylation for both YCCEL1 and SNU719 along with increased gene expression (Figure 5C,D). Interestingly, the magnitude of re-expression was EBV-specific. Consistent with this, we did not see a difference in *RASSF1* expression in tumor samples defined by EBV status, but there was a significantly lower expression of *HOPX* in EBV^+^ tumors (Figure 5E).

**Figure 5.**
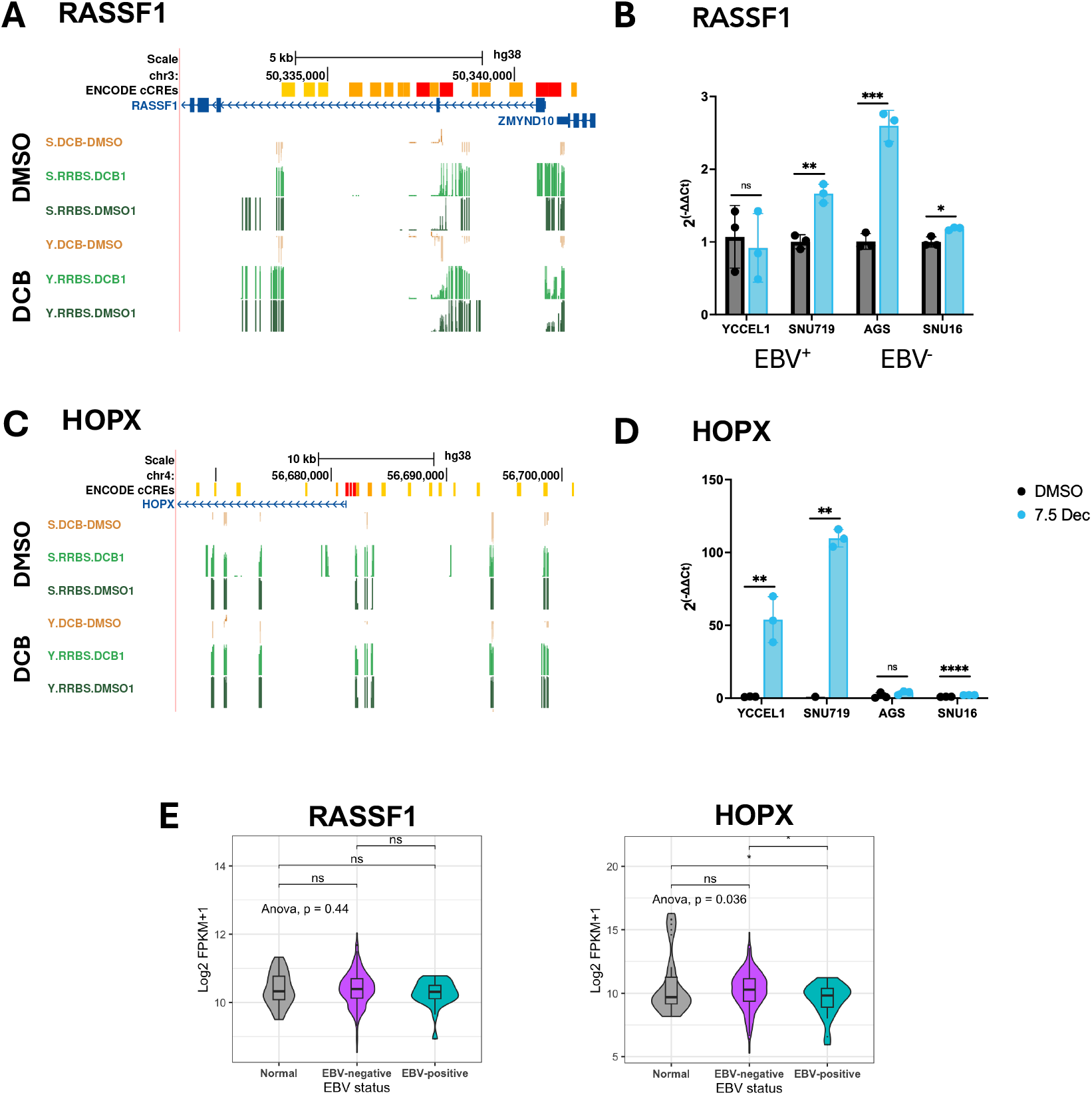
Promoter demethylation correlates with increased gene expression after DCB treatment at key tumor suppressor genes. A) Genome tracks at the RASSF1 gene loci for % change in 5mC after DCB (yellow), % 5mC DCB treated sample (light green), and % 5mC DMSO sample (dark green) for SNU719 (top) and YCCEL1 (bottom). B) Bar plot for expression changes measured by RT-qPCR for RASSF1 after 7d DCB treatment in the gastric cancer cell lines. C) Genome tracks at the *HOPX* gene loci for % 5mC. D) Bar plot of expression changes by RT-qPCR in the gastric cancer cell lines. E) TCGA-STAD cohort expression for RASSF1 and HOPX in normal adjacent tissue (grey), EBV^-^ tumor (purple), and EBV^+^ tumor (blue).

### Integration of RNA-seq and RRBS reveals inverse correlation between promoter demethylation and gene activation in EBV+gastric cancer cells

To better understand the changes in promoter hypomethylation and gene expression, we integrated the RNA-seq and RRBS in the EBV^+^ cell lines and found an inverse correlation between changes in promoter 5mC and transcriptional fold change in both YCCEL1 and SNU719 (Figure A,B). The density coloring of the scatter plots indicate that many promoters did not exhibit a change in 5mC nor a change in expression. In SNU719, 758 genes showed correlated changes in promoter 5mC and transcription compared to 88 in YCCEL1. Of these changes, 55 genes were commonly changed between both cell lines with a significant overlap. Most of these genes have hypermethylated promoters (>80%) before treatment and all showed an increase in expression after DCB (Figure 6D). IPA analysis of concordant genes revealed several pathways common to both YCCEL1 and SNU719, including several signaling pathways including those involved in neuronal signaling and WNT/*μ*-catenin (Figure 6E). Potential upstream regulators identified decitabine, vitamin-a metabolite bexarotene, SMARCA4, SMAD2, POU4F1, FGF8, and ESR2. These analyses highlight epigenetically altered pathways by DCB.

**Figure 6.**
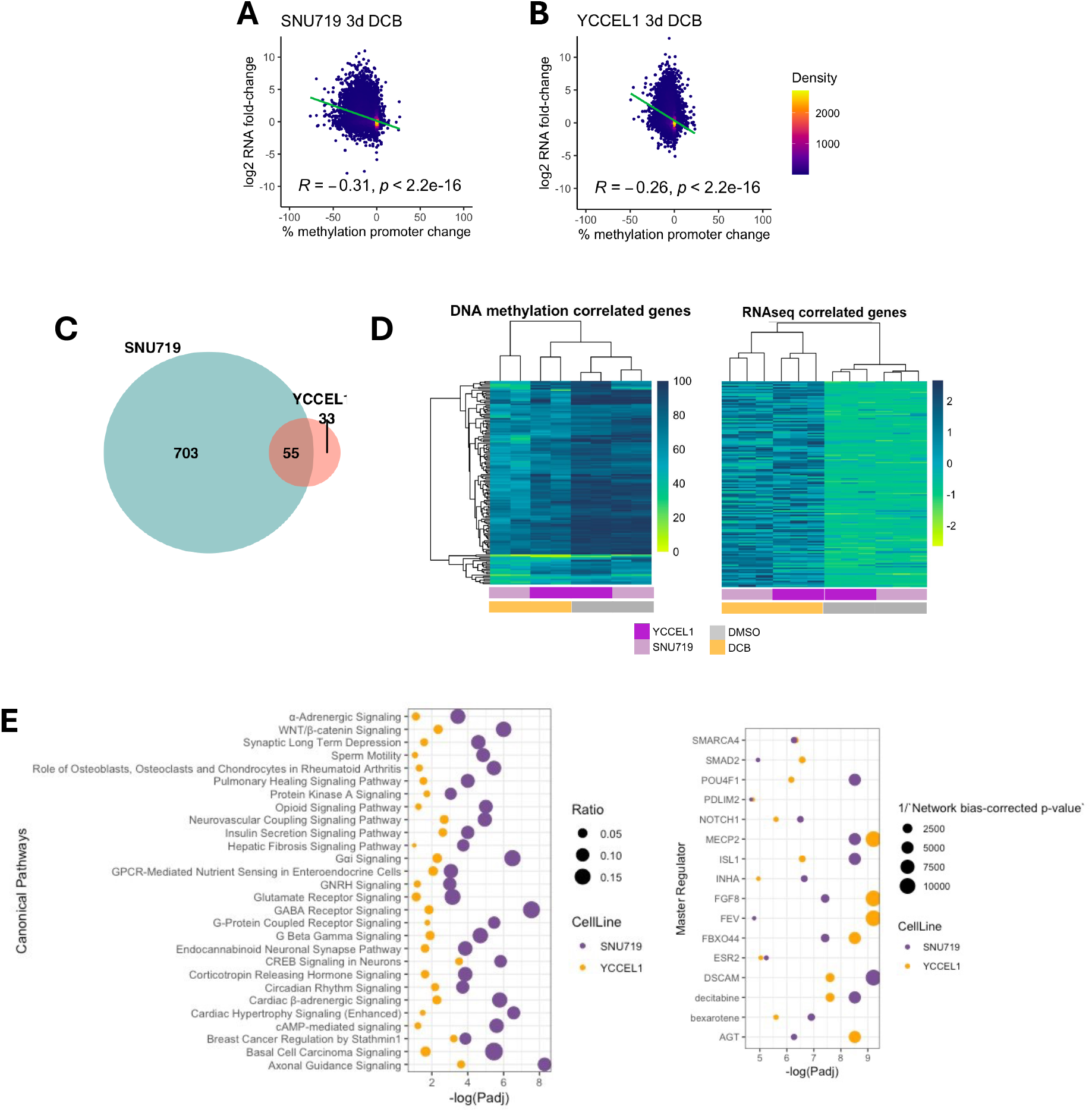
Global analysis of promoter demethylation correlates with increased gene expression after DCB treatment. A) Correlation plot between the effect of promoter methylation change in region 1000bp upstream of the transcriptional start site and the effect on changes in gene expression after DCB treatment for SNU719 and B) YCCEL1. C) Venn diagram showing genes that have greater than 20% loss of promoter 5mC and gene expression log_2_FoldChange > 2 and a significant overlap of 55 genes (p<10-22 hypergeometric test). D) Heatmap of promoter 5mC (left) and gene expression (right) the 55 genes with concordant changes in promoter 5mC and gene expression. E) IPA analysis of genes with concordant changes in promoter 5mC and gene expression for YCCEL1 (n=88, yellow) and SNU719 (n=758, purple) showing common canonical pathways (left) and master regulators (right).

### DCB-induced demethylation enriches RARα motifs and sensitizes EBV+gastric cancer cells to retinoic acid–mediated cytotoxicity and ROS production

To understand regulatory changes on a more global scale outside of the promoter, we performed motif analysis on 50bp regions centered on differentially methylated CpG sites. We identified several known transcription-factor binding sites within these regions (Figure 7A,B). However, the sequence motif of the top 20 transcription factors do not contain CpG dinucleotides suggesting that it is not direct methylation of the DNA excluding binding. Instead, loss of 5mC may indicate a general opening of the DNA allowing access or effecting binding of interacting transcription factor partners (Figure 7C). Furthermore, only a few of the top twenty transcription factors identified are expressed among the gastric cancer cell lines and do not appear to be greatly affected by DCB treatment, including *RARA, ATF1, THRA*, and *MEF2A* (Figure 7D).

**Figure 7.**
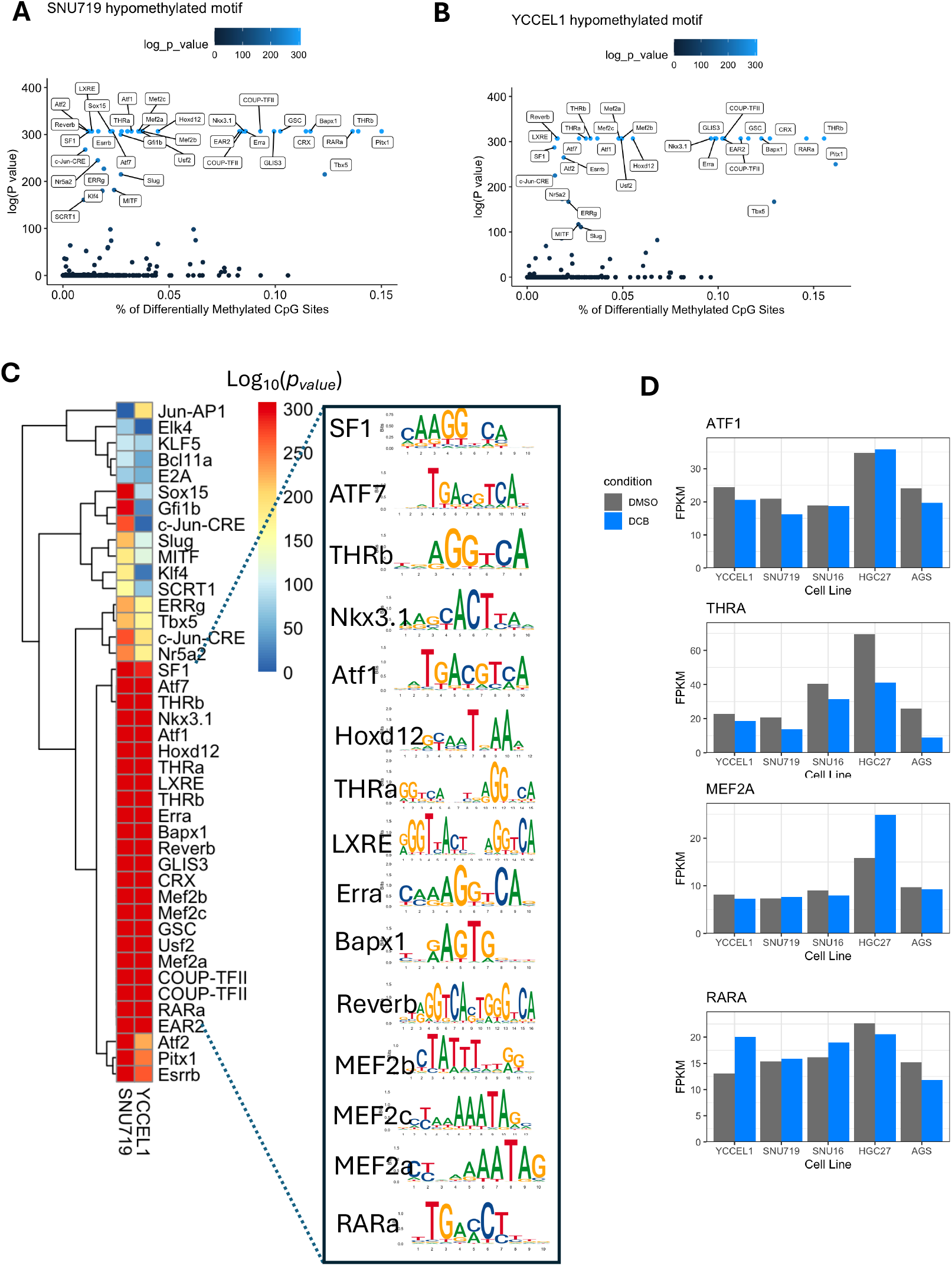
Global analysis of promoter demethylation correlates with increased gene expression after DCB treatment. A) HOMER analysis of known transcription factor motifs enriched in 50bp windows centered on hypomethylated CpG sites for the YCCEL1 cells and B) SNU719. Fraction of CpG sites containing the motif on the x-axis plotted by significance. C) Heatmap of significance displaying the top 20 commonly significant motifs between SNU719 and YCCEL1 with the associated transcription factor logo. RNA-seq FPKM values transcription factors expressed in the gastric cancer cells.

Since retinoic acid metabolite bexarotene appeared as an upstream regulator of transcriptional control and the RAR*α* motif was enriched at demethylated CpG sites and expressed in gastric cancer cells, we looked further into the retinoic acid pathway. *RARB* is a target of RAR*α* signaling and we found a correlation between promoter demethylation and increased gene expression after DCB treatment (Figure 1 A,B). Interestingly, while *RARA* increased expression in most of the cells, *RARB* appeared EBV-specific. Pretreatment of EBV^+^ YCCEL1 cells with DCB to induces epigenetic changes followed by 3d treatment with ATRA showed a significant decrease in live cells compared to single-agent treatment at 3d as measured by PI/Annexin V staining (Figure 8C). Using an antagonist of RAR*α*, AR7, we observed a significant increase in live cells at 6d after DCB treatment. The effect of cellular proliferation was halted by ATRA or in combination with DCB treatment (Figure 8D). Further, we looked at the production of ROS production since it has been previously shown to be generated in DCB and ATRA combination treatment. We found both ATRA and DCB, alone or in combination produced ROS (Figure 8E). We found a transcriptional increase in several genes involved in the production and clearance of ROS after DCB treatment, many regulated by RAR*α* signaling including ROS generating enzymes *DUOX1/2* and *NOX5* as well as antioxidant genes that have been shown to be disrupted by RAR*α* signaling, *GPX1* and *DUSP1* (Figure 8F).

**Figure 8.**
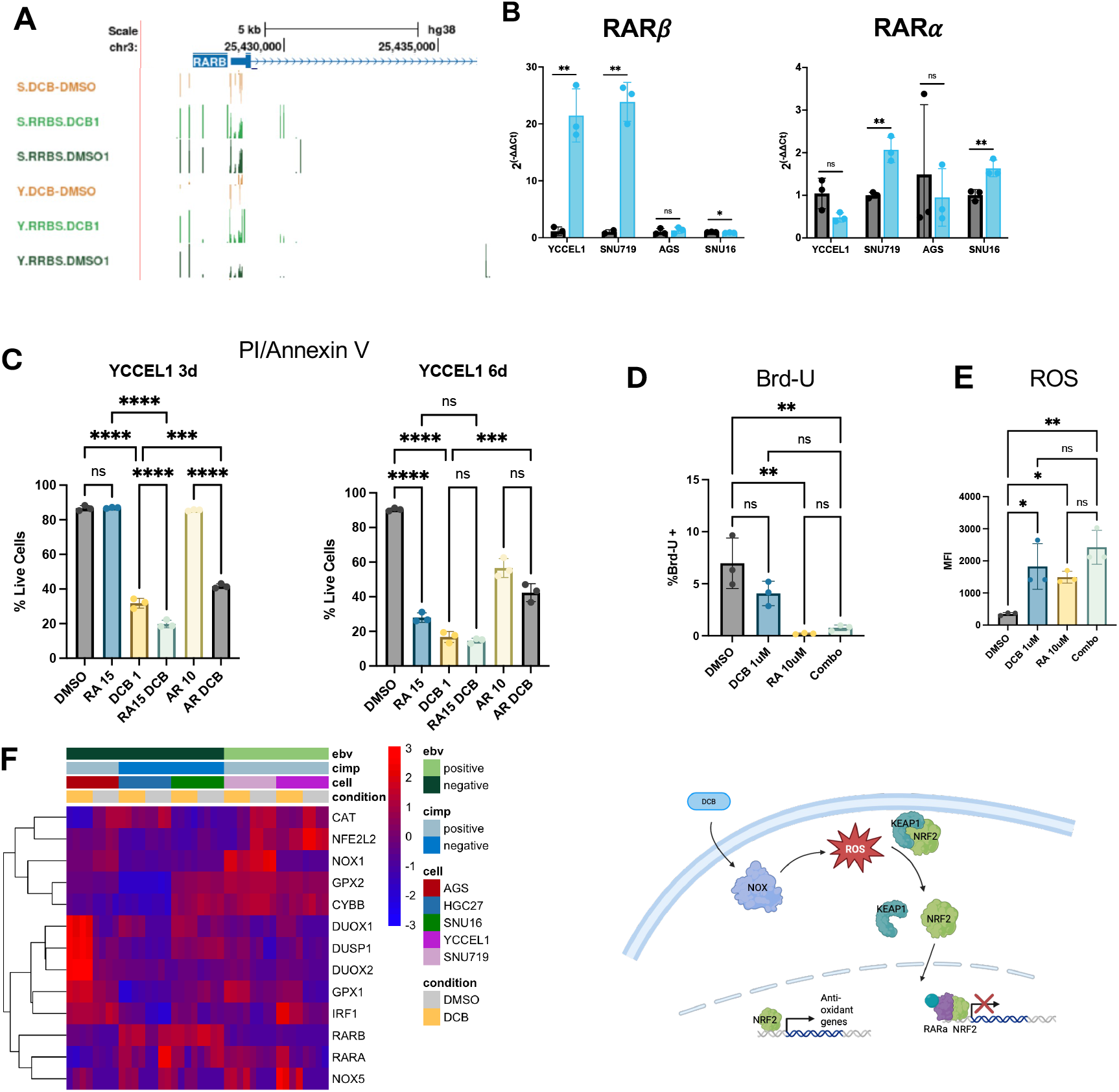
DCB transcriptional effects specific to EBV^+^ cell lines. A) UCSC browser display of RRBS data around the RARB gene. B) Barplots of RT-qPCR data for RARA and RARB genes after DCB treatment. C) PI/Annexin V staining of YCCEL1 cells treated with DCB, tretinoin (RA), and/or RARA antagonist AR7 for three or six days after a 3d pre-treatment of DCB only. D) Flow of 12h Brd-U incorporation after 3d treatment of DCB and/or RA. E) Flow analysis of reactive oxygen species detection (ROS) after 3d treatment with DCB and/or RA. F) Heatmap of RNA-seq data showing genes related to DCB effect on RARA, NRF2/NFE2L2 and ROS production.

## Discussion

Decitabine induced broad epigenetic remodeling in gastric cancer cells, with pronounced effects in EBV^+^ and CIMP-high cell lines. Transcriptomic profiling revealed extensive upregulation of previously repressed genes, many of which are enriched in pathways related to cell differentiation, immune signaling and cancer, and apoptotic signaling being enriched in CIMP positive cell lines. The strong overlap in differentially expressed genes between YCCEL1 and SNU719 highlights a shared EBV-specific transcriptional program, consistent with the viral contribution to CIMP and epigenetic dysregulation in EBVaGC. Importantly, the EBV-specific reactivated genes showed low-basal expression in EBVaGC patients within TCGA, underscoring their potential clinical relevance.

Although global demethylation was widespread, the most robust changes occurred at highly methylated intergenic CpG sites leaving open the possibility of epigenetic manipulation of enhancers within these regions. Promoter regions instead were resistant to 5mC changes in response to DCB, however large changes occurred in some important gene promoters. Integration of RNA-seq and RRBS data confirmed an inverse relationship between promoter 5mC and gene expression, in support of a mechanism where DCB exerts antitumor effects through promoter demethylation and activation of tumor suppressors. In two tumor suppressor genes RASSF1A [52] and HOPX we observed loss of promoter methylation [11, 30, 53]. For RASSF1 we observed loss of promoter methylation in SNU719 and an increase in gene expression. However, in YCCEL1 showed minimal changes in promoter methylation and gene transcription. HOPX showed loss of promoter methylation and YCCEL1 and SNU719 and increased expression after DCB treatment. This appeared EBV specific as these transcriptional changes did not appear in the EBV-cell lines.

Pathway analysis of genes with 5mC promoter changes and motif analyses of regions surrounding differentially methylated CpG sites converged on the retinoic acid receptor (RAR) signaling pathway. The enrichment of RARa motifs around demethylated CpG sites, along with identification of retinoid pathway components as upstream regulators, suggests that DCB induced demethylation enhances accessibility of RARE binding sites and may restore retinoid responsiveness in EBVaGC. Given that EBV-mediated LMP1 activity has been shown to promote RARb hypermethylation and retinoid resistance, these finding imply that DNMTi may counteract viral suppression of differentiation signals. This highlights a potential mechanistic basis for therapeutic interaction between DNMTi and retinoids. These results align with prior evidence in acute myeloid leukemia, where decitabine and all-trans retinoic acid (ATRA) exhibit cooperative antitumor activity through inhibition Nrf2/NFE2L2 induced antioxidant response leading to accumulation of ROS production and cytotoxicity. In line with this, we find large accumulation in ROS by ATRA and DCB. Timing of treatment should also be considered, since ATRA induced cell cycle arrest which may dampen the demethylation effect of DNMTi which depend on mitosis as a mechanism for DNA demethylation. Consistent with this, use of a RAR*α* antagonist and inhibitor of RAR*α* signaling, AR7, showed a rescue in cell death when cells were treated with DCB for 6d. Consistent with this, DNMTi upregulated DUOX1/2 and NOX, enzymes which produce ROS, particularly in the EBVaGC. ROS production activates NRF2 and enables transcription of antioxidant genes such as GPX1 and DUSP1, which we observe upregulated in our EBVaGC. Re-expression of RAR*β* and increased expression of RAR*α* activated by RA can interfere with NRF2 mediated transcription blunting the antioxidant response which aids in cell survival. In this way, DCB can activate the retinoic acid pathway which in turn can increase the toxicity of DCB.

Future studies should systematically map RAR*α* and RAR*β* binding dynamics following decitabine treatment to confirm direct chromatin remodeling at retinoid response elements and if these combinations provide durable cell cycle arrest, apoptosis, or differentiation of EBVaGC. Understanding how EBV factors may be involved in the EBV specific response will be crucial in understanding potential biomarkers and developing EBV specific treatment combinations with decitabine. Particularly EBNA1, which has been shown to effect host-gene transcription to drive cancer progression and small-molecule inhibition shows EBV specificity [33, 34]. Docking sites of the EBV episome are dictated by the host epigenome and it is unclear the impact of decitabine on these interactions and the impact of any resulting changes in host-viral chromatin interactions on host gene expression [19, 54]. In summary, our findings reveal that decitabine induces extensive DNA demethylation in EBV-positive gastric cancer cells, leading to transcriptional reactivation of multiple tumor-suppressive and differentiation-associated pathways such as ATRA. The integrated multi-omics approach to identify reactivation of retinoid signaling provides a compelling rationale for exploring DNMTi combination therapies as a strategy to exploit the unique vulnerabilities of EBVaGC.

## Acknowledgements

We thank all the members of the Tempera laboratory for their support. We are grateful to the Wistar Institute’s Genomics Facility and Bioinformatics Facility for the technical support. Support for the Wistar Institute’s core facilities was provided by Cancer Center Support Grant P30 CA010815. I.T. is supported by R01 AI30209, P01 CA269043, and P30 CA010815. P.M.L., S.S.S. and C.S. are supported by RO1 DE017336, R01 CA259171, R01 CA09360, R01 AI1535086, P01 CA269043, and P30 CA010815. B.E.G. is supported by R01 AI164709, CA228700, P01 CA269043, and U01 CA275301. A.K. is supported by R50 CA211199. S.P.A. is supported by T32 CA09171. Y.W. is supported by American Cancer society grant PF-24-1194768-01-TBE.

